# Identification of genomic signatures and multiple lineage markers from the second and third wave samples of COVID-19 in Western Rajasthan, India

**DOI:** 10.1101/2022.12.10.518819

**Authors:** Naveen Prakash Bokolia, Ravisekhar Gadepalli

**Affiliations:** Viral Research and Diagnostic Laboratory, Microbiology Department, All India Institute of Medical Sciences, Jodhpur (342001), India

**Keywords:** SARS-CoV-2, Genomic signatures, Mutations, Spike domain, Multiple lineage markers, Recombination, Delta and Omicron Variant

## Abstract

Most of the mutations occurred in SARS-CoV-2 are either relatively neutral or swiftly purged. However, some mutations have altered the functional aspects in terms of infectivity and transmission, host-viral interactions, disease severity and immune or vaccine escape. There are emerging evidence that certain mutations are jeopardizing the immune based therapies. The present research report is focused on the identification of genomic signatures of SARS-CoV-2 variant that caused mortality during second and third wave of COVID-19 in Western Rajasthan, India. We identified that Delta clade of SARS-CoV-2 is the predominant cause of mortality during second wave and even third wave in Western Rajasthan, India. Importantly, this study also revealed the unique and common substitution mutations within the spike domain, those are present in mortality and survived persons during the second and third wave of COVID-19 in India. In addition, this study also revealed the multiple lineage markers (Delta and Omicron), that would update with insightful understanding in the clade development of SARS-CoV-2.

## Introduction

Coronaviruses are single stranded positive sense RNA viruses belong to Coronaviradeae family that caused COVID-19 disease (Kim et al., 2020; Zhu et al., 2020). The ongoing COVID-19 pandemic poses one of the greatest global threats in modern history and has already caused severe social and economic costs. The development of efficient and rapid sequencing methods to reconstruct the genomic sequence of SARS-CoV-2, the etiological agent of COVID-19, has been fundamental for the design of diagnostic molecular tests and to devise effective measures and strategies to mitigate the diffusion of the pandemic (Aleem et al., 2021; Banho et al., 2021; Ignatieva et al., 2022). Whole genome sequencing data of SARS-CoV-2 has important role in the elucidation of lineage of virus (with evolution), tracing the pathogenicity, transmission and spread within a particular state or local region (Ignatieva et al., 2022; Kannan et al., 2021; Lu et al., 2020; Naveca et al., 2021; Tegally et al., 2020; Washington et al., 2021; Zhan et al., 2020).

According to one CDC report (April, 2022), Omicron variant infection occurs among the 10 persons who already have had infection with Delta variant within 90 days. Delta variant remains predominant followed by overtaken by Omicron variant (Roskosky et al., 2022). In India, similar scenario happened, where the Omicron associated infections risen during Jan-Feb 2022.

Rajasthan state shares 5.66% population of total Indian population, and Jodhpur district has second largest population (after Jaipur) with the estimate of 4,276,374. Thus the present study is designed to provide a broad overview of SARS-CoV-2 variant specification that is/was circulated among the survived persons and hospitalized patients who died during the hospitalization at tertiary care unit of All India Institute of Medical Sciences, Jodhpur, India. Although, during third wave the number of deaths were comparatively very less in India (Arnaout and Arnaout, 2022; Jha et al., 2022). However, it is important to determine the prime cause of infection with morbidity and mortality in a specified region, and to update the information about previous and current SARS-CoV-2 variants in circulation.

Within this perspective we categorized the SARS-CoV-2 samples in four different categories. First category involves “second wave survived”, second category involves “second wave hospitalized and deceased”, third category involves “third wave survived” and fourth category involves “third wave hospitalized and deceased”. The comparative analysis (between different categories) of genomic signatures was done with respect to spike domain of SARS-CoV-2. The present study revealed that Delta is the causative agent of mortality in third wave of COVID-19 as well, and not the Omicron variant of SARS-CoV-2. In addition, the multiple lineages are detected from the second wave samples of COVID-19, those might be important in determining the genomic aspects of clade development of SARS-CoV-2. This study updates and defines the unique and common mutations in the RNA genome of SARS-CoV-2 variants.

## Materials and Methods

### Categorization of SARS-CoV-2 positive samples used in this study

This study was designed to identify SARS-CoV-2 variants and genomic signatures from the second and third wave of COVID-19 patients those were died during hospitalization at tertiary care unit of AIIMS Hospital, Jodhpur, Rajasthan. This study does not involve specific parameters those are based on gender, age, sex or associated disease complications. In the present investigation already stored samples during a particular COVID-19 wave (peak phase) was used, however only samples those fit in to the particular Ct value criteria (RNA integrity) were processed further for sequencing. We involved 50 SARS-CoV-2 positive samples those were collected from non-hospitalized individuals, 50 samples from hospitalized and died patients during second wave of COVID-19. Similarly, we involved 35 SARS-CoV-2 positive samples those were collected during third wave of COVID-19, and 16 samples from the patients those died during hospitalization at tertiary care unit of AIIMS Jodhpur, Rajasthan. We could involve only 16 samples from third wave mortality because of the lower cases of hospitalization and mortality during third wave. In addition, a parallel comparison was done with SARS-CoV-2 genome sequencing data obtained from the samples of non-hospitalized (survived) COVID-19 positive individuals.

### Real Time RT-PCR of SARS-CoV-2 positive samples

The stored SARS-CoV-2 positive samples (−80°C storage) were taken and properly thawed at room temperature. The total RNA of samples was extracted by using QIAamp Viral RNA kit. In order to proceed for sequencing experiments, we determined the Ct value of extracted RNA samples by real time RT-PCR. The positive samples those fall within the range of 20-25 Ct values were further proceeded for whole genome sequencing.

### Whole genome sequencing of SARS-CoV-2 on MiSeq system (Illumina platform)

The whole genome sequencing of SARS-CoV-2 was performed on MiSeq System (Illumina technology), that is installed at R’VRDL, Microbiology Department, AIIMS Jodhpur. The library preparation was done by COVIDSeq Assay Kit and the protocol was followed according to manufacturer’s protocol instructions. The library was quantified on Qubit fluorimeter and 12 Pico molar concentration of final library was loaded for sequencing run. MiSeq Reagent Kit v3 (150-cycle) was used for sequencing run after final library loading. After the completion of sequencing run sequencing data was obtained in FASTQ format and subsequently used for analysis.

### Analysis of sequencing data for clade verification, mutational profiling, genomic signatures and sharing the data at GISAID

The initial analysis was performed by using the Illumina^®^ DRAGEN COVID Lineage App. These mutations were listed in Table 2. Subsequently FASTA files were used for mutational analysis, and was done on CoVsurver mutations app-GISAID that delivers total number and types of mutations in spike domain of SARS-CoV-2. The clade analysis (assignment), verification, and phylogenetic tree was constructed by Nextclade v1.14.1 software. The sequencing data of all the samples relevant to this study has been shared on GISAID and accession IDs have been provided in supplementary data file (Khare et al., 2021).

**Table 1:** Table represent the number of samples belong to particular lineage of SARS-CoV-2, during the second and third wave of COVID-19.

|  | Different category of SARS-CoV-2 samples used in the present investigation |  |  |  |
| --- | --- | --- | --- | --- |
| Type of SARS-CoV-2 lineage | Second Wave Non-Hospitalized (May 2021) | Second Wave Hospitalized Patients with mortality (May-June 2021) | Third Wave Non-Hospitalized Individuals (March-May 2022) | Third Wave Hospitalized Patients with mortality (March-May 2022) |
|  | Total number of samples used and respective numbers belong to particular lineage |  |  |  |
|  | 50 | 50 | 35 | 16 |
| B.1.1 | 1 | 3 | Not Detected | Not Detected |
| B.1.617.1 | 1 | 4 | Not Detected | 1 |
| B.1.617.2 | 21 | 21 | Not Detected | 7 |
| AY.122 | 20 | 14 | Not Detected | 7 |
| AY.38 | 4 | 4 | Not Detected | Not Detected |
| AY.46 | Not Detected | 1 | Not Detected |  |
| **BA.2** | Not Detected | Not Detected | 9 | Not Detected |
| **BA.1.1**<br>**BA.1.17**<br>**BA.1.18**<br>**(Omicron:**<br>**BA.1-like)** | Not Detected | Not Detected | 26 | Not Detected |
| **Unassigned**<br>**(Omicron)** | 3 | 3 | Not Detected | 1 |
| **Unassigned**<br>**(Delta)** | Not Detected | Not Detected | Not Detected | Not Detected |

**Table 2:** The table represents the total number of mutations present in spike domain within each categorized set of samples.

| **Lineage** | **Total Representative mutations in spike domain of SARS-CoV-2 during second wave (associated with survived COVID-19 positive individuals)** | **Total Representative mutations in spike domain of SARS-CoV-2 during second wave (associated with hospitalization and mortality)** | **Total representative mutations in spike domain of SARS-CoV-2 during third wave (associated with hospitalization and mortality)** |
| --- | --- | --- | --- |
| **AY.122**<br>**Lineage** | **T19R E156G F157del R158del G446V L452R T478K D614G P681R D950N** | **T19R R21I A67V H69del V70del T95I <u>G142D</u> V143del Y144del Y145del E156G F157del R158del T307I L452R T478K D614G Q675H P681R(674) D950N N969K** | **T19R ins108AV <u>G142D</u> E156G F157del R158del N185D L452R T478K D614G P681R D950N** |
| **B.1.617.2 Lineage** | **T19R A67V H69del V70del T95I E132D E156G F157del R158del A222V G446V L452R T478K E583D D614G H655Y N679K P681H P681R A845S N856K D950N Q954H N969K L981F** | **T19R T29A A67V H69del V70del K77T T95I S98F G142D W152L E156G F157del R158del V213G A222V L452R T478K Q613H D614G P681R N764K D796Y D950N N969K L981F A1070S** | **T19R A67V H69del V70del K77T T95I E156G F157del R158del V213G A222V L452R T478K D614G P681R D950N N969K** |

### Phylogenetic analysis of multiple lineage variants

In this analysis, we involved six FASTA sequences of multiple lineage variants, two FASTA sequences of B.1.617.2 lineage, two FASTA sequences of AY.122 lineage and two FASTA sequences of B.1 lineage. The FASTA files of multiple lineage variants were analysed by using NGPhylogeny.fr tool. The analysis was done in “one-click’’ mode(Lemoine et al., 2019). The resultant phylogenetic tree was exported, and presented as Figure 1.

**Figure 1:**
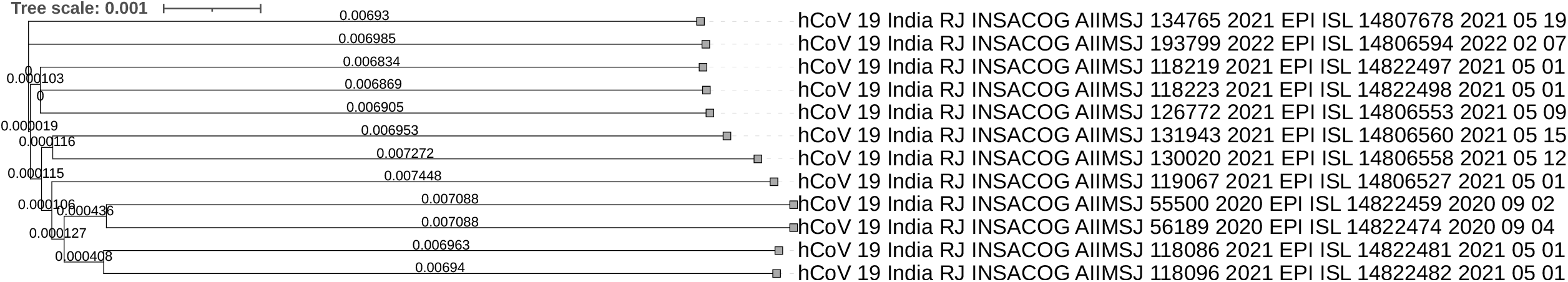
The presented phylogenetic tree indicates the possibility of evolution of mixed lineages due to intervariant recombination events between the RNA genome of SARS-CoV-2. The analysis also includes FASTA sequences of first wave and second wave variants. The GISAID accession IDs of first wave variants are EPI_ISL_14822459 and EPI_ISL_14822474. The GISAID accession IDs of second wave variants (lineage B.1.617.2 and AY.122) are EPI_ISL_14822481, EPI_ISL_14822482, EPI_ISL_14822497 and EPI_ISL_14822498 respectively. Remaining six sequences belong to multiple lineages. The phylogenetic tree indicates that RNA genome of B.1.617.2 lineage and B.1 lineage undergoes recombination events, subsequently multiple recombination events could have occurred that leads to development of SARS-CoV-2 clade.

## Results

### B.1.617.2 and AY.122 lineages (Delta clade) remain persistent during third wave of COVID-19 in Western Rajasthan

In the present study report we investigated second wave (Delta wave), third wave and post third wave SARS-CoV-2 positive samples. Samples from respective wave was divided in to non-hospitalized and hospitalized (with mortality) category, and comparative variant analysis was done. We involved 16 SARS-CoV-2 positive samples (available) in the category of “hospitalized and died during third wave”. Among them, in 14 samples we detected Delta and Delta plus variant that is categorized in B.1.617.2 and AY.122 lineages respectively. Remaining two patients were found to be infected with Kappa (B.1.617.1) and unassigned Omicron respectively. Sequencing output from the “third wave mortality patients” revealed that B.1.617.2 and AY.122 lineages of Delta clade remain the most persistent variants of SARS-CoV-2 during the third wave of COVID-19. In addition, study also revealed that Delta variant of SARS-CoV-2 still remain the causative agent of death instead of Omicron variant that has been characterized during Jan-2022 to March-2022 (third Wave in India). Apart from mortality cases, the sequencing results from non-hospitalized COVID-19 individuals (third wave) revealed the Omicron variant of BA.1.1 and BA.2 lineages mainly (Table 3). The present research report indicates that during third wave of Omicron Clade, 14 SARS-CoV-2 patients died due to Delta variant instead of Omicron variant (Table 1).

**Table 3:** Table presents the total number of substitution mutations present in a particular lineage during third wave of COVID-19.

|  |  |  |
| --- | --- | --- |
|  | Total Representative mutations in spike domain of SARS-CoV-2 during third wave (COVID-19 positive) associated with BA.2.38 Variant | Total Representative mutations in spike domain of SARS-CoV-2 during third wave (COVID-19 positive) associated with BA.2 Variant |
| Mutations in Spike domain | **T19I L24del P25del P26del A27S G142D V213G G339D S371F S373P S375F T376A D405N R408S K417N K417T Q498R N501Y Y505H N556K D614G V642G H655Y N679K P681H N764K D796Y Q954H N969K** | **T19I L24del P25del P26del A27S L84I G142D N185D V213G Y248N G339D R346T S371F S373P S375F T376A D405N R408S K417N N440K G446S L452M N460K S477N T478K E484A Q493R Q498R N501Y Y505H D614G H655Y T676A N679K P681H N764K D796Y Q954H N969K** |

Several studies have suggested that previously infected individuals with earlier circulating SARS-CoV-2 may have reduced protection from reinfection with the B.1.351 variant.

### Identification of persistent or stable mutations within Delta clade during post COVID-19 phase

The Variant of Concern (VOC), Delta, was first reported in India in December 2020, and was responsible for the deadly second wave in April 2021. It was then reported in USA in March 2021, and became the most dominant variant there, and had been detected throughout the world including UK. Studies reported that specific mutations in the spike domain makes the properties of virus deleterious in terms of infectivity, disease severity and pathogenicity. In this regard, there are several mutations those remain conserved during different waves of COVID-19 with respect to particular variant of SARS-CoV-2. These three categories were put in to the context of analysis because most of the samples belong to Delta clade only. It can be easily observed that the number of mutations decreased with the time in Delta clade. Even though the patients were died in third wave due to infection from the B.1.617.2 lineage and AY.122 lineage, however, the number of mutations in spike domain is significantly lower in comparison to second wave patient samples (including both mortality and survived). It indicates that the core mutations in spike domain remain invariable in a manner where the RNA biology of virus does not change in terms of infectivity and pathogenicity. These core mutations could be further helpful in the development of particular drug or vaccine, while considering the mutations in Delta clade in current scenario as well.

The comparative analysis between “second wave non-hospitalized”, “second wave Hospitalized (mortality)” and “third wave Hospitalized (mortality)” leads to identification of unique amino-acid substitution mutations within the spike domain of SARS-CoV-2. We characterized these mutations as “unique substitution mutations in spike domain” in the Delta clade, as these are unique in the group of mortality and survived patients of second wave and third wave in India.

Present study revealed that Delta variant of SARS-CoV-2 remained dominant in terms of pathogenicity and infectivity irrespective of ongoing emerging variants, specifically in Western Rajasthan, India.

### Identification of multiple lineages from the second wave samples of COVID-19

The present study involved sequencing data analysis of 100 SARS-CoV-2 positive samples from the second wave of COVID-19 in India (2021), those belonged to deceased and survived patients. Out of 100 samples, sequencing results from the eight samples revealed the mixed mutations from the both Delta and Omicron variants (Table 4). These submissions are marked with *“ML’’* and belong to multiple lineages. The identified (listed) mutations are highlighted in bold in Table 4. These mutations were found to be present in the spike domain. Apparently, some of the Omicron specific mutations have been known/studied for the potential impact on SARS-CoV-2 pathogenesis. Within this context, Omicron specific mutations N679K and H655Y are present at the interface/junction of the S1 and S2 subunit, and both are associated with increased infectivity. In addition, deletion mutations at N-terminal domain (H69-V70) are associated with increased infectivity, antibody escape and diagnostic escape.

**Table 4:**
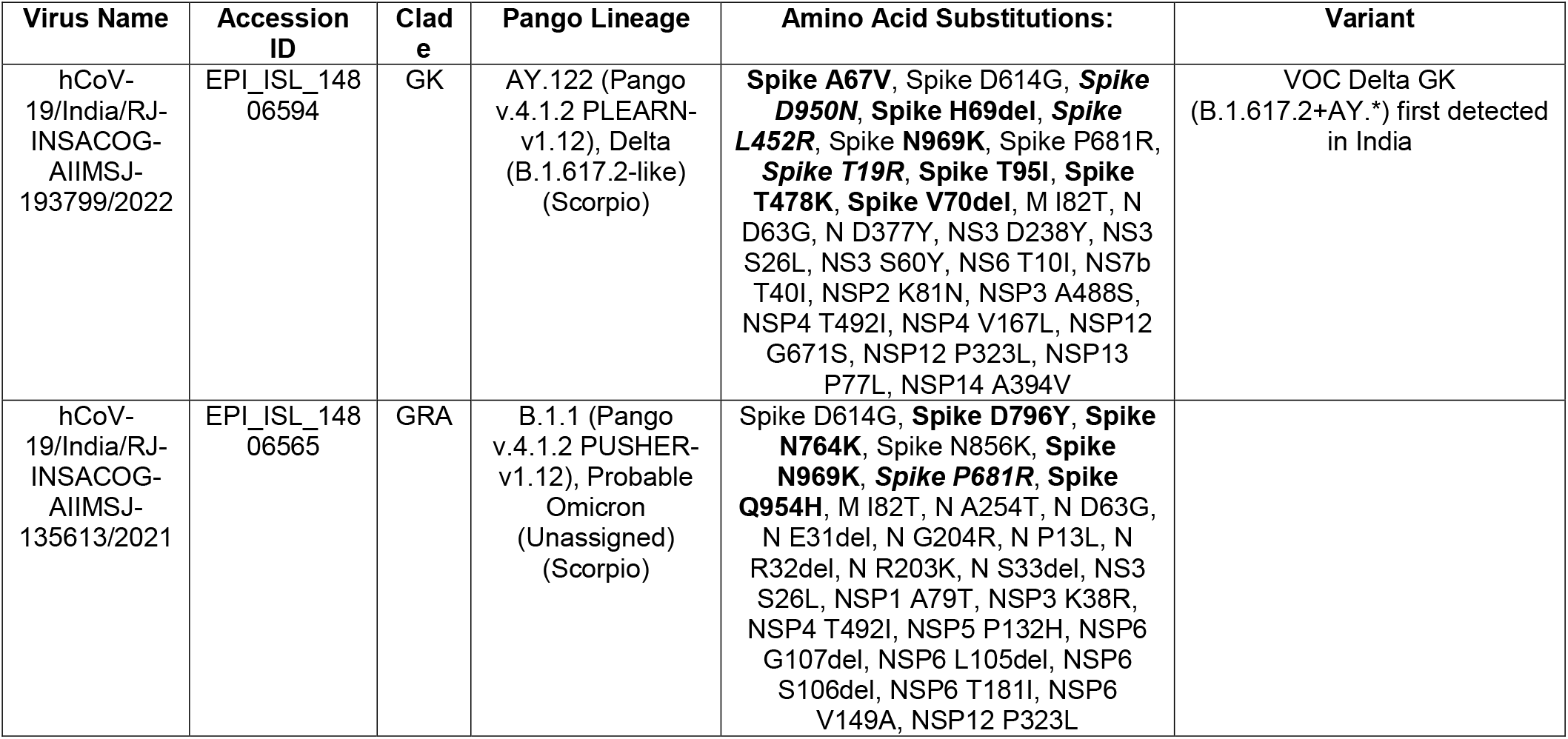
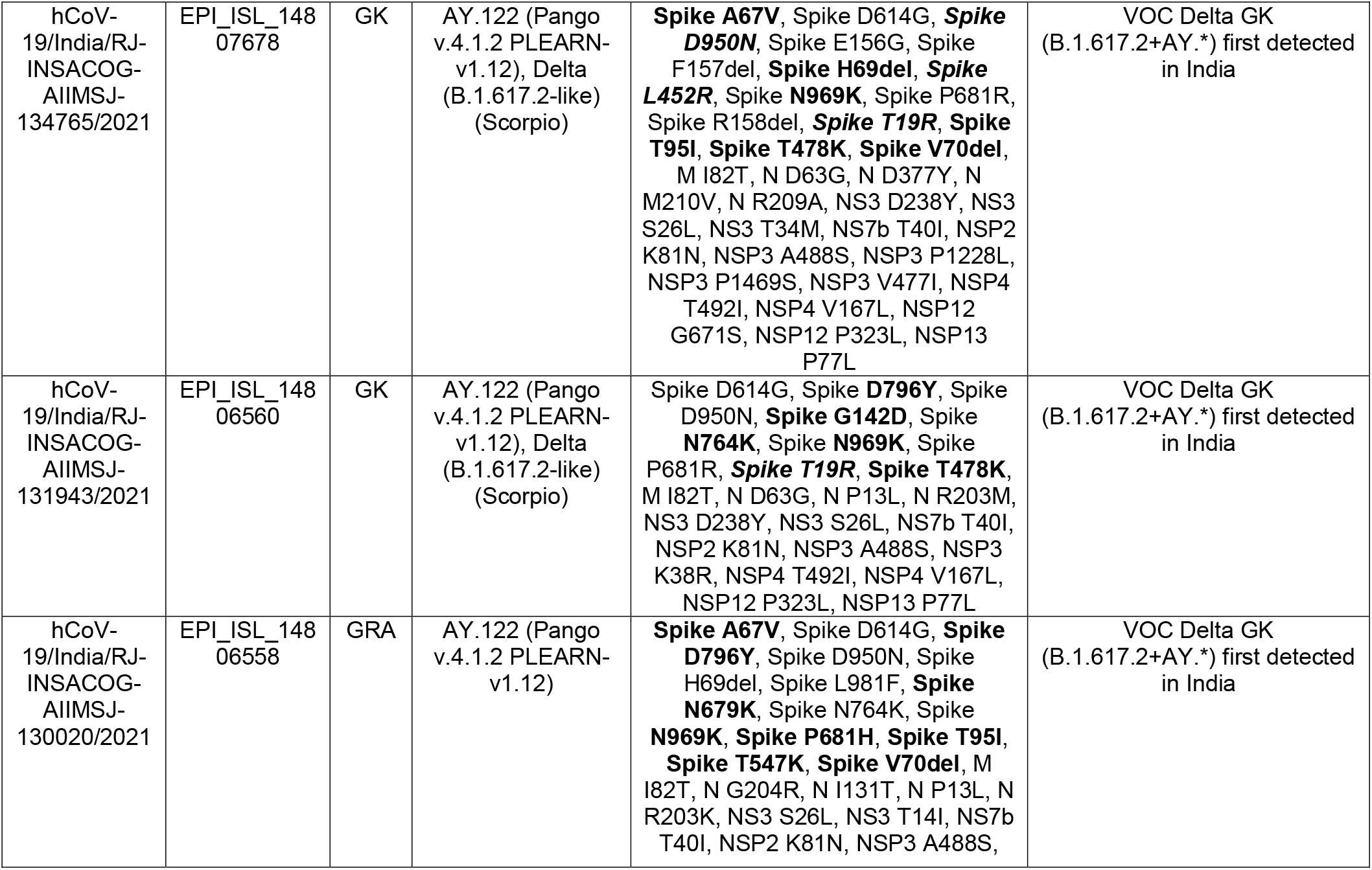
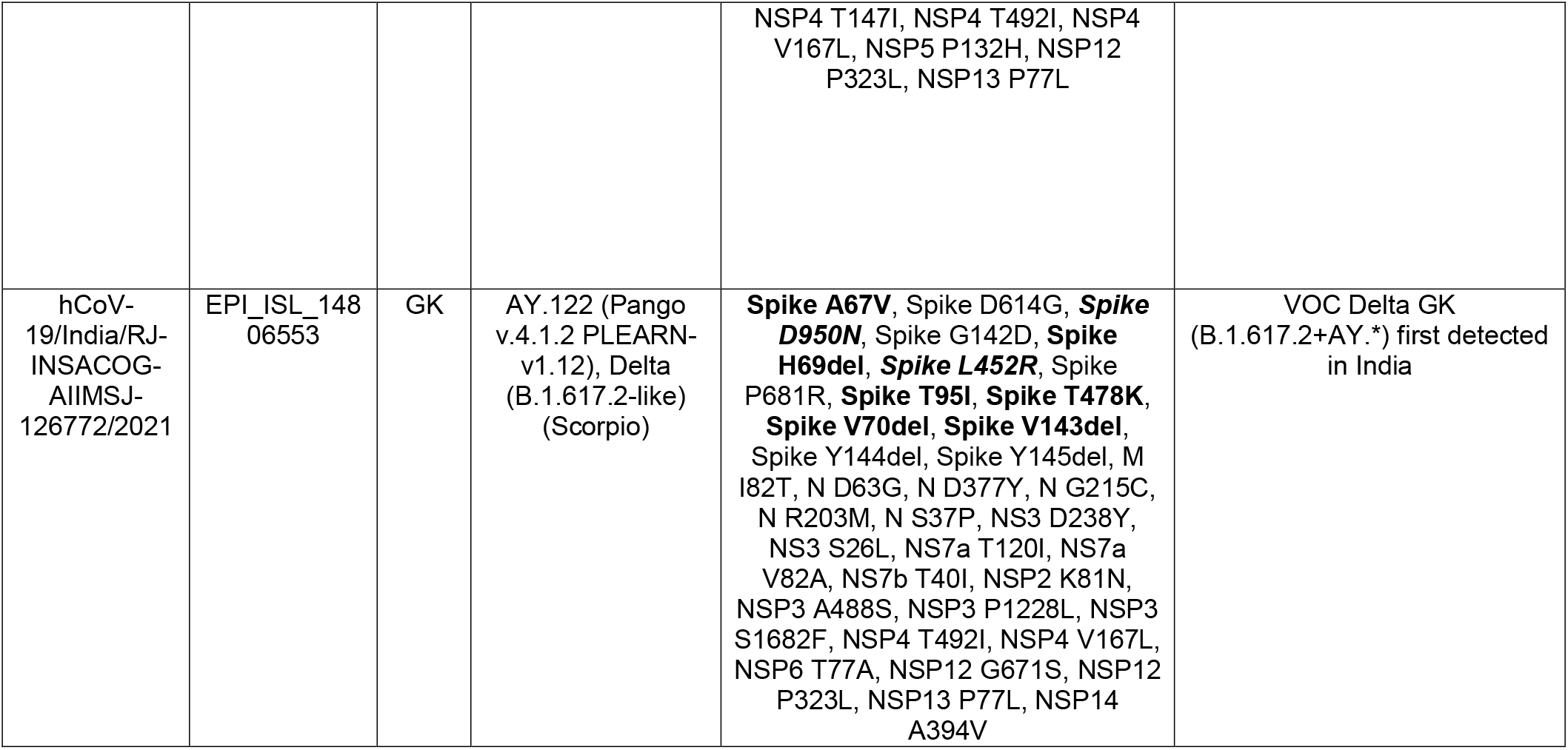
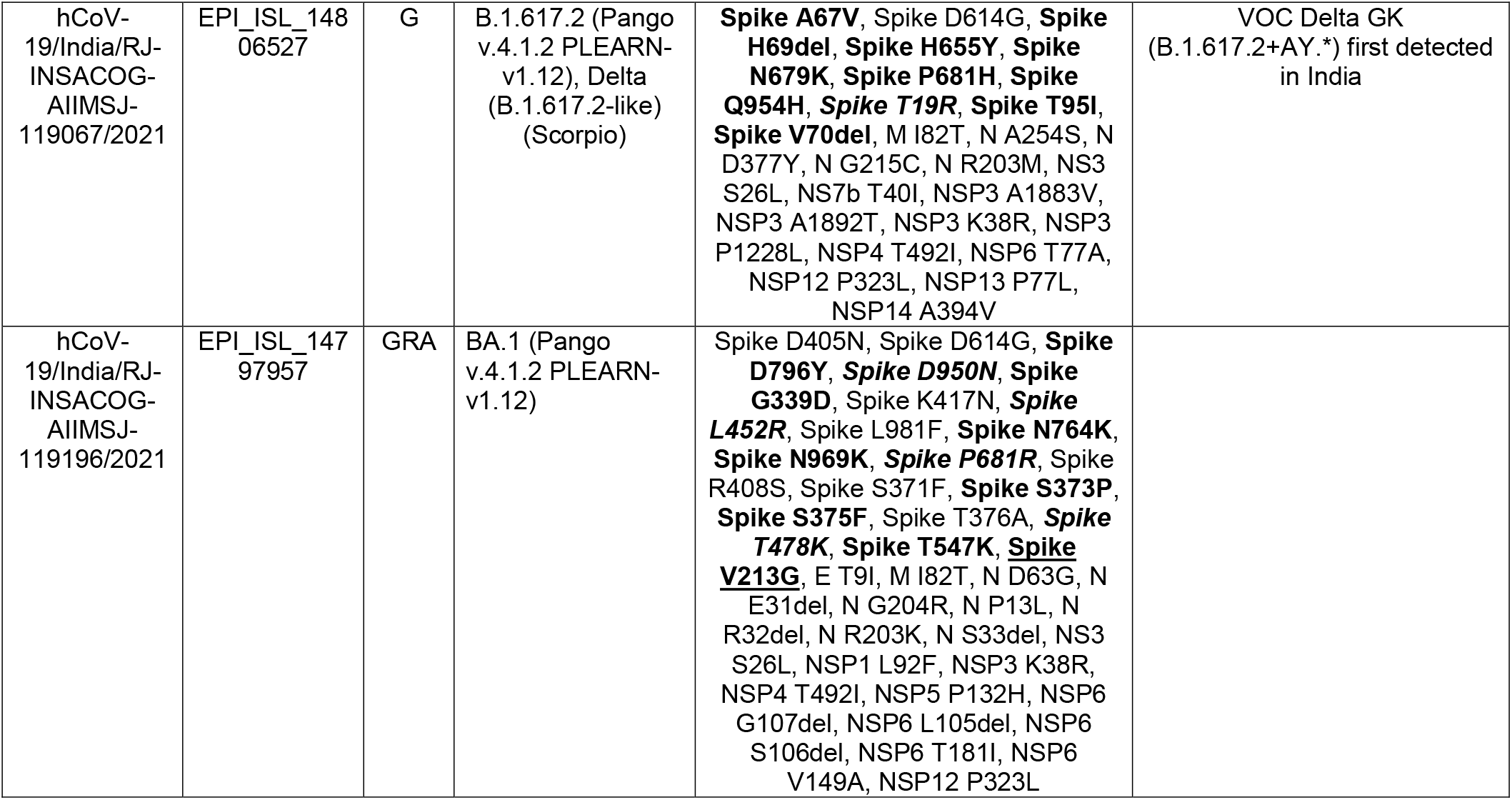
Table presents all of the mutations those are present in mixed lineages and unassigned (probable) Omicron variant during third wave of COVID-19 in India. Defining mutations of Delta clade are presented in bold only, whereas defining mutations of Omicron clade are highlighted with bold and italics.

Importantly, mutations in receptor binding domain have also been found in one mixed lineage (EPI_ISL_14797957) and was characterized as BA.1 Pango lineage. This lineage bears some of the defining mutations of Delta variant in spike domain: L452R, P681R and T478K, however other major mutations like T19R and 156del and 157del mutations were not present. Same mixed lineage also bears Omicron specific substitutions: G339D, S373P, S375F those could be involved in increased infectivity, higher immune escape and transmissibility (Alkhatib et al., 2022; Cao et al., 2022; Liu et al., 2022; Long et al., 2020; Ou et al., 2022). Previous research reports mentioned about the G339D substitution mutation as rare mutation in SARS-CoV-2 genome database before the detection of Omicron, and this mutation is associated with significant increase in binding affinity with the ACE2 receptor (Alkhatib et al., 2022; Starr et al., 2020). Other than these, the significance of spike domain mutations like A67V and G142D in relation to the possible impact of SARS-CoV-2 pathogenesis has not been demonstrated.

In order to get insights on the evolution pattern of mixed lineages, we analysed the sequences of mixed lineages along with B.1, B.617.2 and AY.122 lineages. Since, the mixed lineage samples derived from the second wave of COVID-19, therefore, we purposefully involved first wave and second wave FASTA sequences during the analysis of phylogenetic tree. The mixed lineage indicates the possible recombination events between the GK and G clade variants (Figure 1). Subsequently, these events could have occurred with the RNA genome of different SARS-CoV-2 variants. The hierarchy pattern of mixed lineage SARS-CoV-2 clade development indicates that amino acid substitutions might have been started during the second wave (Delta wave) of COVID-19 in India that later on established as distinct Omicron clade.

## Discussion

The fourth VOC, Delta, was first reported in India in December 2020, and was responsible for the deadly second wave of COVID-19 (Jha et al., 2022). It was then reported in USA in March 2021, and became the most dominant variant there, and had spread throughout the world including UK. The lineage is comprised of three subtypes: B.1.617.1, B.1.617.2 and B.1.617.3, with diverse mutations in NTD and RBD of spike protein with higher immune evasion potentials. The B.1.617.2 and B.1.617.3 variants differ from each other mainly by two mutations: T478K and E484Q in the RBD. The Delta variant was described more transmissible than the viruses causing SARS, MERS and Ebola as well as influenza and common cold, and as contagious as chickenpox by CDC (https://www.yalemedicine.org/news/5-things-to-know-delta-variant-covid). It was reported that there was the rise in number of cases (10 fold) and mortality (3 fold) in India, with the emergence of Delta variant, displacing alpha and kappa lineages (Dhar et al., 2021). A Scottish study demonstrated that the risk of hospitalisation was higher (double) in Delta infections as compared to Alpha, in non-vaccinated groups (Sheikh et al., 2021). These are combining perspectives that inform about the underlying reason of persistent impact of Delta clade that remains during third wave of COVID-19 as well. In present study, this impact was identified in terms of mortality in COVID-19 patients during the hospitalization at tertiary care unit in Western Rajasthan, India. Whereas Omicron variant was reported to be of lower risk in terms of serious disease, hospitalization and mortality in India and other countries (South Africa, England, Scotland, Denmark and Canada) (Espenhain et al., 2021; Singhal, 2022; Wolter et al., 2021).

In addition, present study identifies the unique amino acid substitution mutations in the mortality group of patients in Delta clade, that could be of further consideration and helpful for future vaccine development. For example, the unique amino acid substitution mutations can be used in a cumulative manner or in an independent manner, according to further docking or experimental outcome that results in increase in affinity or any significant observation. In addition, the reduction in number of mutations in spike domain is described among the group of “third wave mortality with hospitalization”. These markedly reduced number of mutations in spike domain further signifies that RNA biology of SARS-CoV-2 remains unchanged depending on the type and respective context of mutation, however a significant number of core mutations (that defines a particular Clade or lineage of variant) must be present in order to propagate at same pathogenicity and infectivity.

This study revealed that mixed lineage bearing the amino acid substitutions (as markers) from the both Delta and Omicron variants. The role of amino acid substitutions has been identified in the Omicron specific variant that modulates the potential impact of SARS-CoV-2 pathogenesis (Alkhatib et al., 2022; Ou et al., 2022). These findings indicates the possibility that during the infection cycle, the RNA genome of SARS-CoV-2 may have undergone through the spontaneous mutations and deletions events. In addition, the second probability could be of recombination events between the RNA genome of GK and G clade variants of SARS-CoV-2. The probability of recombination events between Alpha and Delta variants could be higher because the mixed lineages have the amino acids substitutions from the both Delta and Omicron variants. Since Omicron variant has more mutations then Alpha, Beta, Gamma and Delta, those promotes stronger immune escape and accelerated transmission (Xu et al., 2022). The accumulation of increased mutations signify the impact and possibility of recombination events. Some of the recent research reports have provided the evidence of possible intervariant and intravariant recombination of Delta and Omicron variants (Focosi and Maggi, 2022; Ou et al., 2022; VanInsberghe et al., 2021). One study provided the clear evidence of iterlineage recombination events between B.1.1.7 and B.1.177 lineages, where recombinant variant had spike gene from B.1.1.7 variant (Jackson et al., 2021). In addition, recent studies point out towards the human chronic infections and/ or animal reservoirs could be the contributing factor in the evolution of RNA genome of SARS-CoV-2 (Tegally et al., 2022). This study signifies the identification of mixed lineage markers from the second wave of COVID-19 in India. These markers (amino acid substitutions) would be insightful and significant in understanding the evolution pattern of the SARS-CoV-2 RNA genome within the context of recombination events.

## Supporting information

Supplementary Data Files 1 and 2.zip

## Data Availability

All the sequencing data relevant to article, has been submitted on GISAID (Khare et al., 2021). The supplementary file of respective sequences (accession IDs) has been made available as supplementary data.

## Acknowledgment

We are very grateful and thankful for becoming the part of INSACOG. We are very grateful to NIV Pune, as a mentor institute of AIIMS Jodhpur, and providing the sequencing kits for the whole genome sequencing of SARS-CoV-2.

